# Identification of a link between splicing and endoplasmic reticulum proteostasis

**DOI:** 10.1101/2023.09.28.559974

**Authors:** Muhammad Zahoor, Yanchen Dong, Marco Preussner, Sabrina Shameen Alam, Renata Hajdu, Veronika Reiterer, Stephan Geley, Valerie Cormier-Daire, Florian Heyd, Loydie A. Jerome-Majewska, Hesso Farhan

**Author notes:** MZ and YD contributed equally to this work.

## Abstract

The role of general splicing in endoplasmic reticulum (ER)-proteostasis remains poorly understood. Here, we identify SNRPB, a component of the spliceosome, as a novel regulator of export from the ER. Mechanistically, SNRPB regulates the splicing of components of the ER export machinery, including Sec16A, a regulator of ER exit sites. Loss of function of SNRPB is causally linked to cerebro-costo-mandibular syndrome (CCMS), a genetic disease characterized by bone defects. We show that heterozygous deletion of SNRPB in mice resulted in intracellular accumulation of type-1 collagen as well as bone defects reminiscent of CCMS. Silencing SNRPB inhibited osteogenesis in vitro, which could be rescued by overexpression of Sec16A. This indicates that the role of SNRPB in osteogenesis is linked to its effects on ER export. Finally, we show that SNRPB is a target for the unfolded protein response (UPR), which supports a mechanistic link between the spliceosome and ER-proteostasis. Our work highlights SNRPB as a novel node in the proteostasis network, shedding light on CCMS pathophysiology.

## Introduction

Proteostasis refers to the intricate balance between protein synthesis, trafficking, and degradation (Klaips *et al*, 2018; Plate & Wiseman, 2017). The endoplasmic reticulum (ER) plays a crucial role in proteostasis as it handles one-third of the proteome, making it a major hub for maintaining a balanced proteome. Newly synthesized secretory proteins leave the ER in a COPII-dependent manner at specialized domains referred to as ER exit sites (ERES) (Kurokawa & Nakano, 2019; Peotter *et al*, 2019). ERES represents an important node in the proteostasis network, because they are major determinants of ER unloading. If the function or capacity of ERES does not match the quantity of proteins queuing for export, then the result is an overload of the ER, which will trigger the unfolded protein response (UPR). The UPR involves three transmembrane proteins, IRE1α, PERK, and ATF6α, which possess stress-sensing domains in the ER and signaling effector domains in the cytosol (Walter & Ron, 2011). The objective of the UPR is designed to restore ER homeostasis by increasing the folding capacity of the ER by inducing chaperones (Bakunts *et al*, 2017). In addition, the UPR increases the ER-export capacity by inducing the expression of ERES regulators (Farhan *et al*, 2008). This emphasizes the important role of ERES in ER-proteostasis.

Previous comprehensive screenings have focused on identifying regulators of the secretory pathway, leading to the discovery of novel ERES regulators (Farhan *et al*, 2010; Simpson *et al*, 2012; Wendler *et al*, 2010). However, these screens may have overlooked certain regulators, such as components of the splicing machinery, which are largely absent. Alternative splicing has been shown to be involved in ER export regulation (Wilhelmi *et al*, 2016), but little is known about the role of general splicing in ER proteostasis. Given that the UPR increases mRNA expression of genes encoding ERES components, it is plausible that mRNA splicing contributes to ERES regulation.

To address knowledge gap, we focused on the spliceosomal subunit SNRPB for two main reasons. To investigate this further, we focused on the spliceosomal subunit SNRPB for two reasons. Firstly, SNRPB was identified in a genome-wide screen for factors regulating autophagy, a process closely linked to ERES structure and function (Cui *et al*, 2019; Farhan *et al*, 2017; Zahoor & Farhan, 2018). Secondly, genetic alterations leading to reduced SNRPB expression have been linked to Cerebro-costo-mandibular syndrome (CCMS), a disorder characterized by skeletal malformations (Bacrot *et al*, 2015; Lynch *et al*, 2014). However, the specific role of SNRPB in bone development and its pathophysiological basis remain unclear. While we are aware that a defect in splicing might result bone alterations due to multiple pathways, we think that a defect of ERES function is a strong candidate. Mutations in various components of the COPII machinery or UPR signaling pathway have been associated with bone formation disorders (Boyadjiev *et al*, 2006; El-Gazzar *et al*, 2023; Garbes *et al*, 2015; Horiuchi *et al*, 2016; Zheng *et al*, 2021). This commonality led us to hypothesize a potential link between SNRPB and the ER proteostasis network. SNRPB is a Sm protein that together with other Sm proteins (D1-3, E, F, & G) forms the Sm-ring, which is a core scaffold of the U1, U2, U4, and U5 small ribonuclear proteins (snRNPs) (Schwer *et al*, 2016).

In this work, we show that the spliceosomal component SNRPB regulates the levels of several ERES components, including Sec16A. We found SNRPB to be induced by the UPR signal transducer ATF6, and to thereby mediate ERES biogenesis under conditions of secretory cargo overload. Our work thereby establishes an unprecedented role for mRNA splicing in the regulation of ER proteostasis.

## Results

### SNRPB regulate ER-to-Golgi trafficking

We first determined whether loss of SNRPB affects ER-to-Golgi trafficking using the retention-using-selective-hook (RUSH) technique (Boncompain *et al*, 2012), which monitors the synchronous release of a wave of fluorescently tagged proteins from the ER. We tested two types of RUSH reporters, which were characterized by us and others in the past: Mannosidase-II (Boncompain *et al*., 2012; Phuyal *et al*, 2022) and collagen-X (Stadel *et al*, 2015). Knockdown of SNRPB by two different siRNAs (Figure S1A) affected the trafficking of both RUSH reporters (Figure 1A-B), indicating a possible broad effect on ER export that is not dependent on the type of cargo. Using live imaging mannosidase-II trafficking, we observed that the trafficking defect is a delay in trafficking, rather than a complete block (Figure S1B). To rule out that the observed effects are due to overexpression, or due to the fact that the RUSH system relies on synchronized trafficking waves, we performed an immunofluorescence staining for ERGIC-53, a cargo receptor that cycles between the ER and the ERGIC. The number of peripheral ERGIC-53 positive puncta is dependent on intact export from the ER, and blocking this trafficking step results in reduced ERGIC puncta (Ben-Tekaya *et al*, 2005; Farhan *et al*., 2010). In line with our RUSH experiments, we observed that silencing SNRPB resulted in a reduction in the number of peripheral ERGIC-53 puncta (Figure 1C).

**Figure 1.**
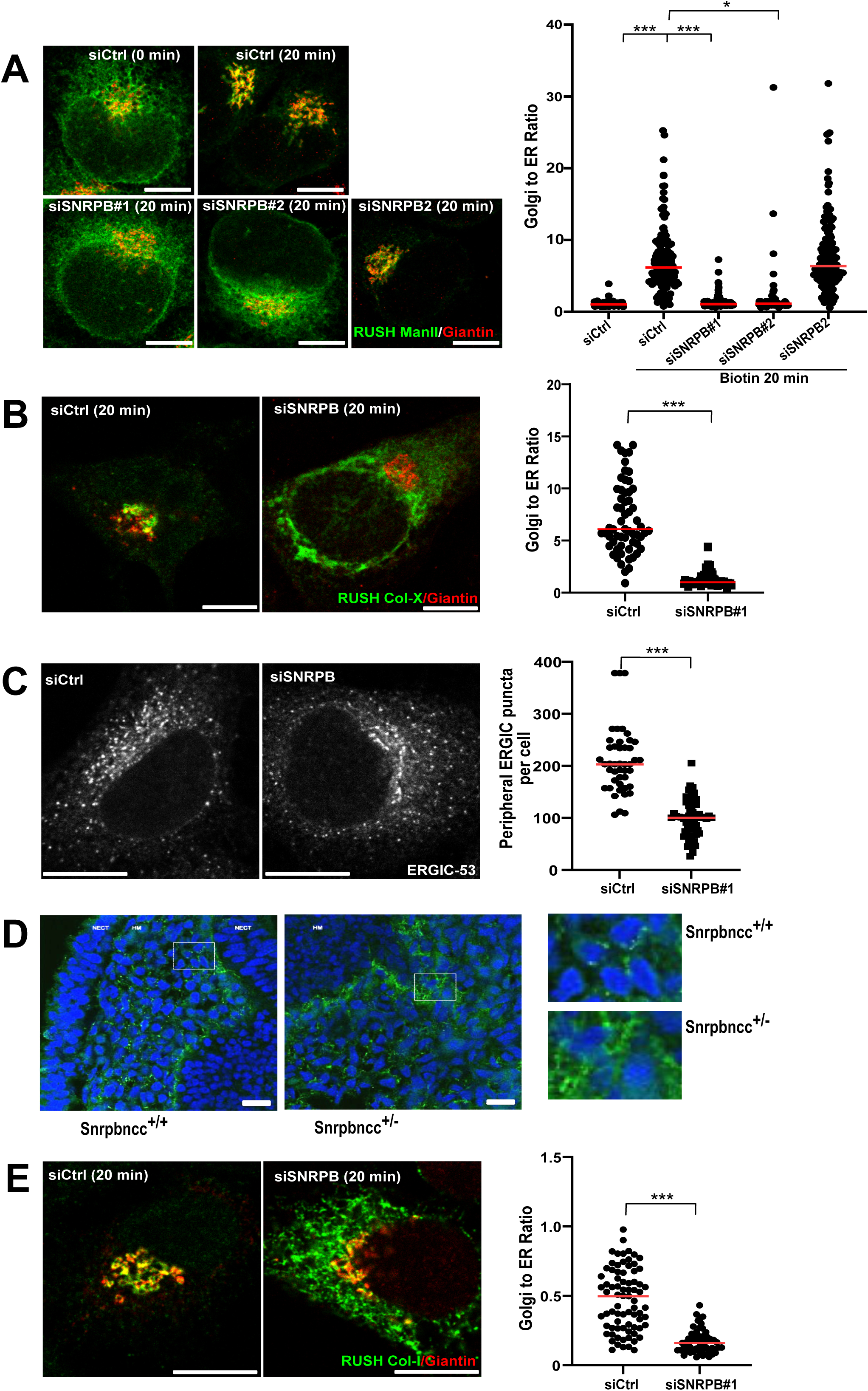
Depletion of SNRPB inhibits ER-to-Golgi trafficking. ***A***, HeLa cells stably expressing mCherry tagged mannosidase-II RUSH construct (ManII) were transfected with siRNA against SNRPB (siSNRPB) or with a non-targeting control siRNA (siCtrl). After 72 h, cells were treated with biotin (40 µM) for 20 min followed by fixation and immunofluorescence staining against giantin to label the Golgi. Graph to the right shows a quantification of 3 independent experiments. The fluorescence intensity of the mCherry signal in the Golgi area was normalized to that in the whole cell giving the Golgi-to-ER ratio. *** indicates a p-value smaller than 0.001 and * indicates a p-value smaller than 0.05, obtained one-way ANOVA. ***B***, HeLa cells were transfected with siRNA against SNRPB (siSNRPB) or with a non-targeting control siRNA (siCtrl). After 48 h, cells were transfected with a plasmids encoding the GFP-tagged type-X collagen as part of the RUSH system. After 24 h, cells were treated with biotin (40 µM) for 20 min, followed by fixation and immunofluorescence staining against giantin to label the Golgi. *** indicates a p-value smaller than 0.001 obtained using a non-paired, two-tailed t-test. ***C***, HeLa cells were transfected with siRNA against SNRPB (siSNRPB) or with a non-targeting control siRNA (siCtrl). After 48 h, cells were fixed and immunostained against ERGIC-53. Graph to the right shows a quantification of 3 independent experiments where the number of ERGIC puncta was counted. *** indicates a p-value smaller than 0.01 obtained using a non-paired, two-tailed t-test. ***D***, immunofluorescence staining of collagen (green) and nuclei (blue) in the head region of E9.5 embryos of *Snrpb* control (*Snrpb^+/+^; Wnt*^tg/+^) and mutant (*Snrpb^ncc/+^*) embryos. ***E***, HeLa cells were transfected with siRNA against SNRPB (siSNRPB) or with a non-targeting control siRNA (siCtrl). After 48 h, cells were transfected with a plasmids encoding the type-I GFP-tagged collagen as part of the RUSH system. After 24 h, cells were treated with biotin (40 µM) for 20 min, followed by fixation and immunofluorescence staining against giantin to label the Golgi. The fluorescence intensity of the collagen signal in the Golgi area was normalized to that in the whole cell. Graph to the right shows a quantification of 3 independent experiments. *** indicates a p-value smaller than 0.001 obtained using a non-paired, two-tailed t-test.

Loss of function of SNRPB underlies the rare genetic disease CCMS, which is characterized by bone abnormalities such as micrognathia and rib defects. In light of the effect of SNRPB on ER export, we hypothesized that loss of SNRPB might cause defects in trafficking of type-I collagen. We therefore performed immunofluorescence staining in neural crest tissue from wild type mice and *Snrpb* neural crest specific heterozygous (*Snrpb^ncc+/-^*), which we described recently(Alam *et al*, 2022). In the tissue of wild type mice, collagen-I was distributed mostly extracellularly and to some extent intracellularly (Figure 1D). However, collagen-I exhibited a different staining pattern in tissue from *Snrpb^ncc+/-^* mice, with a stronger intracellular distribution. This might be indicative of a trafficking defect. To further test this, we used a collagen-I construct that was engineered into the RUSH system(McCaughey *et al*, 2019) and found that silencing SNRPB inhibits the trafficking of collagen-I (Figure 1E).

To gain a further insight into the mechanism of the retardation of ER-to-Golgi transport, we determined the number of ERES. Silencing SNRPB resulted in a marked decrease of ERES (Figure 2A). The effect of SNRPB on ERES was also confirmed in the U2OS cell line (Figure S1D). Depletion of SNRBP2 did not affect ERES number (Figure 2A), which is in line with an absence of an effect on ER-to-Golgi transport. SNRPB is mutated in CCMS patients, and therefore we examined ERES in fibroblasts from a patient with a mutation on 165G>C in CCMS (Bacrot *et al*., 2015). To rescue the expression of this spliceosomal component, we transfected the patient-derived cells with a plasmid encoding GFP-tagged wild type SNRPB, which localized to the nucleus (Figure 2B), similarly to endogenous SNRPB (Figure S2). After 24 h, cells were fixed and immunostained against Sec31 to label ERES. When we compared cells expressing SNRPB with non-transfected cells, we observed a marked increase in the number of ERES (Figure 2B), which is in line with the role of SNRPB in the regulation of ER export. Altogether, these data show that the spliceosomal subunit SNRPB regulates ER-to-Golgi trafficking by affecting ERES number and function.

**Figure 2.**
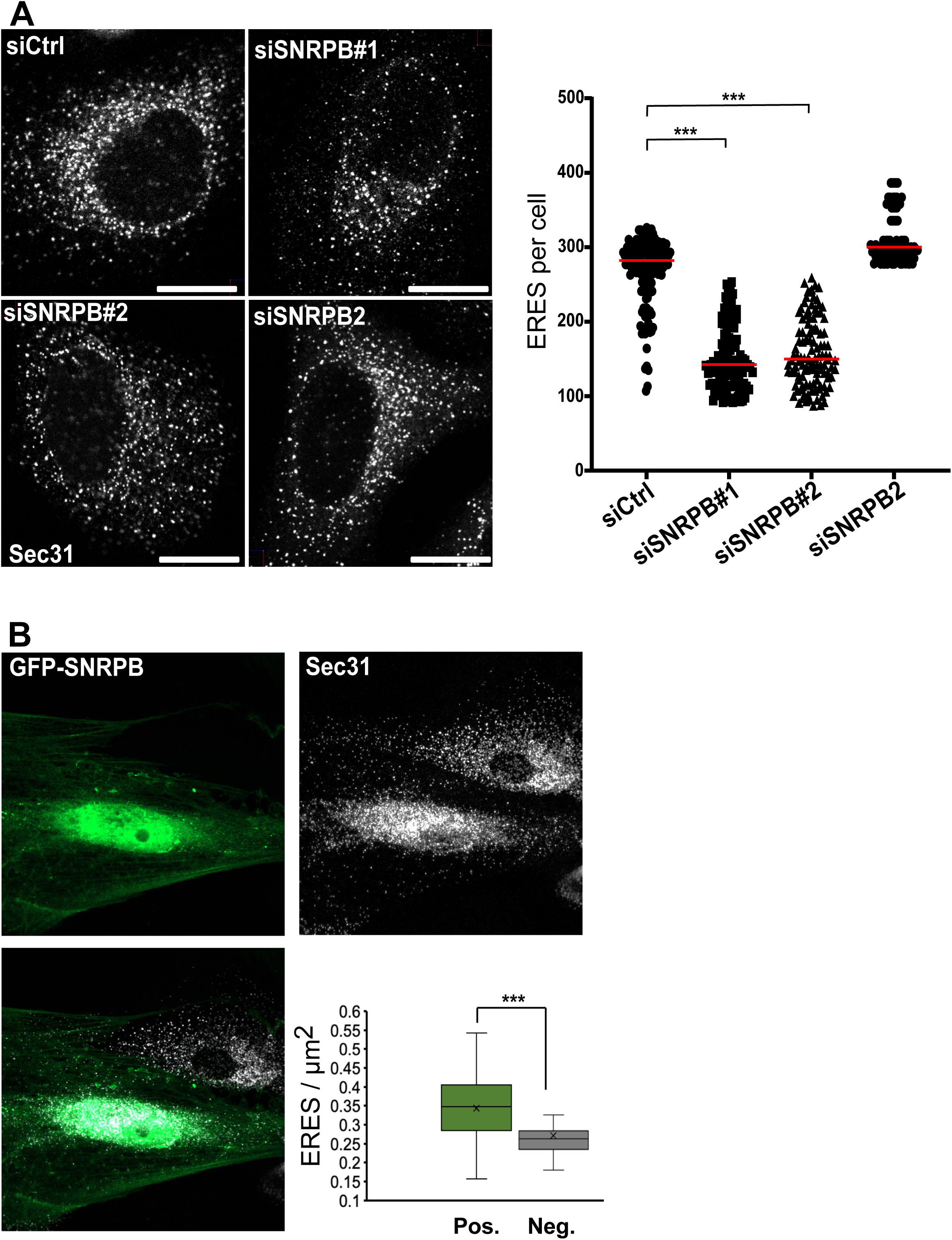
SNRPB regulates the number of ERES. ***A***, HeLa cells were transfected with siRNA against SNRPB or SNRPB2 or with a non-targeting control siRNA (siCtrl). After 72 h, cells were fixed and immunostained against SEC31 to label ERES. The number of ERES per cell was counted using ImageJ and is displayed in the graph to the right. *** indicates a p-value smaller than 0.001 obtained using one-way ANOVA. ***B***, fibroblasts derived from a patient with SNRPB mutation were electroporated with a plasmid cDNA encoding GFP-tagged SNRPB. After 24 h, cells were fixed and immunostained against Sec31 to label ERES. Box blot shows a quantification of a total of 60 cells from 3 independent experiments. *** indicates a p-value smaller than 0.001 obtained using a non-paired, two-tailed t-test.

### ER-to-Golgi trafficking is regulated by Sm-ring components, but not by other SNRPs

SNRPB is part of the Sm-ring together with SNRPD1-3, SNRPE, SNRPF, and SNRPG. We asked whether knockdown of components of the Sm-ring affects the levels of the other subunits. Using qPCR, we found that silencing of SNRPB had no major effect on the mRNA levels of the Sm-ring components SNRPD1, SNRPF and SNRPC (Figure 3A). SNRPG was found to be upregulated, which might be due to a compensatory effect. We also did not find an effect of SNRPB depletion on the levels of SNRPA and SNRPC (Figure 3A), which are unrelated SNRPs, not part of the Sm-ring. Next, we tested whether the effect on ER-to-Golgi trafficking is observed specifically with SNRPB knockdown, or whether any alteration of the Sm-ring would affect secretory trafficking. We therefore silenced the Sm-ring components SNRPB, SNRPD1, and SNRPG and observed that all of them negatively affected ER-to-Golgi trafficking as assayed using a RUSH assay with mannosidase-II as a reporter (Figure 3B). Silencing of the non-related SNRPB2, and SNRPC had no appreciable effect on ER-to-Golgi trafficking (Figure 3B& Figure 1A). These results reinforce the conclusion of a link between the general spliceosome and ERES function.

**Figure 3.**
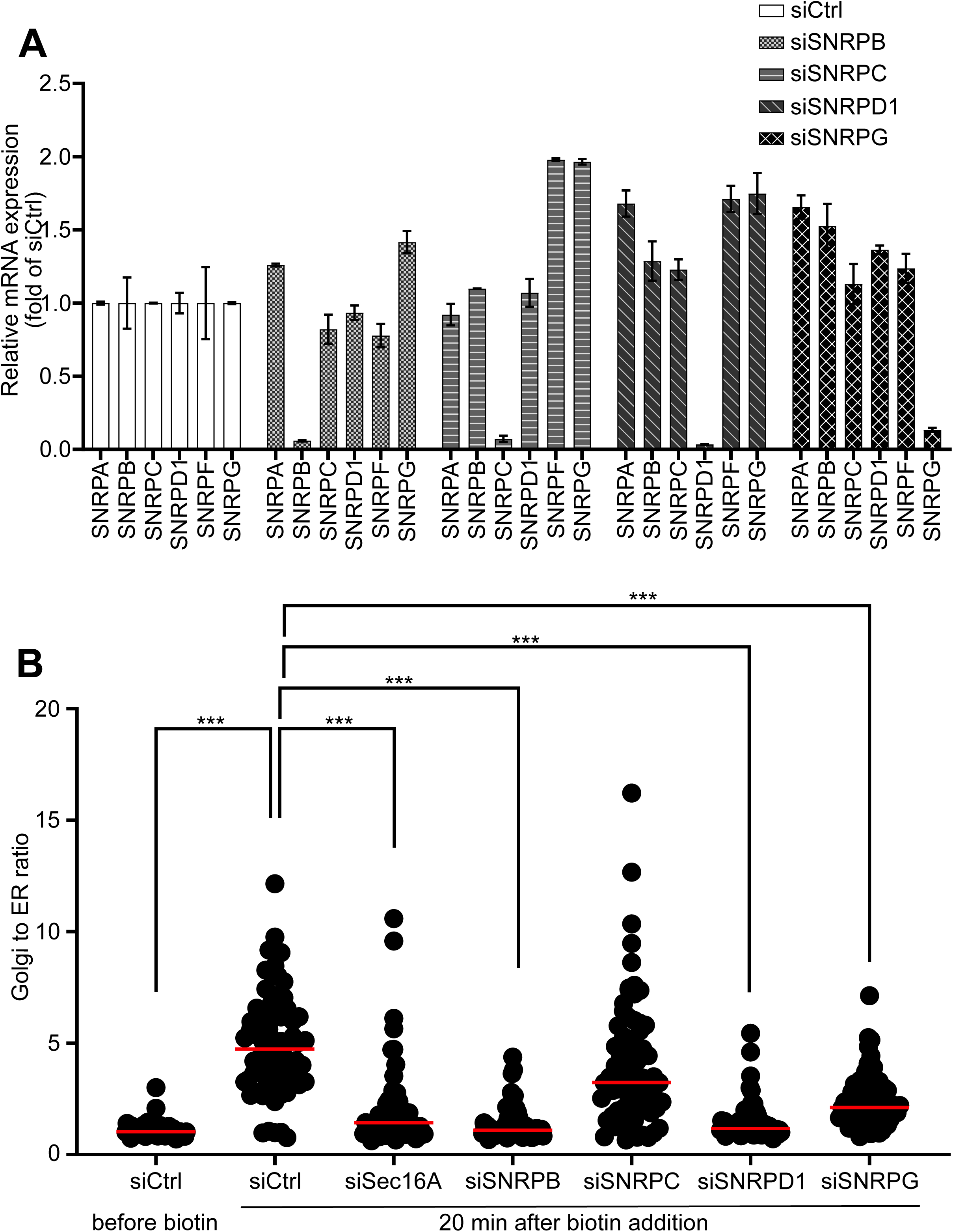
Sm ring components regulate ER-to-Golgi trafficking. ***A***, HeLa cells were transfected with siRNA against the indicated transcripts. siCtrl indicates transfection with non-targeting control siRNA. After 72 h, qPCR was performed to measure the levels of the mRNAs of the Sm-ring components SNRPB, SNRPC, SNRPD1, SNRPF and SNRPG as well as for the non-Sm-ring SNRPBA. ***B***, HeLa cells expressing mannosidase-II RUSH construct were transfected with siRNA as indicated. siCtrl indicates transfection with non-targeting control siRNA. After 72 h, the RUSH assay was started by treatment with 40 μM biotin followed by fixation and immunostaining for GM130 to label the Golgi. The graph shows a quantification of 3 independent experiments. The fluorescence intensity of the signal in the Golgi area was normalized to that in the whole cell giving the Golgi-to-ER ratio. *** indicates a p-value smaller than 0.001 obtained using one-way ANOVA.

### SNRPB knockdown decreases the level of ERES components

We next wanted to explore the cause for the alteration in ERES number and function. To this end, we determined the level of several components of ERES in SNRPB knockdown cells. We noticed a down-regulation of several proteins involved in ER-to-Golgi trafficking such as Sec16A, Sar1A, Sec12, Sec24C, and Sec31 (Figure 4A). There was a weak reduction in the protein levels of Sec13, which we consider to rather be a consequence of the reduction of Sec31A levels, the dimerization partner of Sec13. Two pools exist for Sec13, one that is part of the COPII coat and another one that is part of the nuclear pore complex (Enninga *et al*, 2003). We think that the weak reduction of Sec13 is because only the COPII-associated pool is affected because of a loss of its dimeric partner Sec31. We also probed for the Golgi matrix protein Giantin, but did not find a difference as was the case for the ER chaperone calnexin (Figure 4A). We used actin as a loading control, which was also not affected by the depletion of SNRPB.

**Figure 4.**
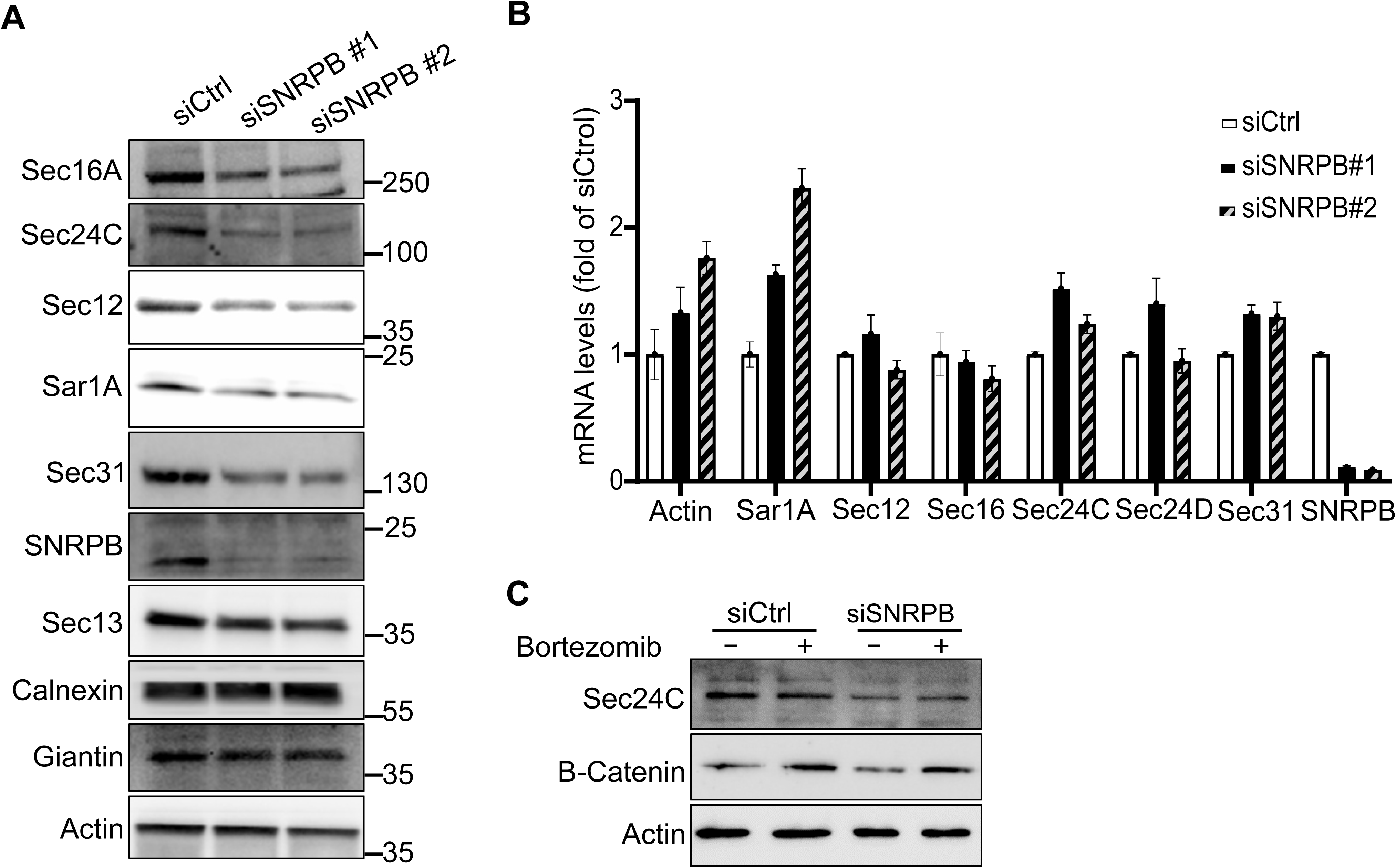
Depletion of SNRPB regulates the protein levels of ERES components. ***A***, HeLa cells were transfected with two siRNAs against SNRPB or with a non-targeting control siRNA (siCtrl). After 72 h, cells were lysed followed by immunoblotting as indicated. ***B***, quantitative RT-PCR of the indicated transcripts in HeLa cells depleted of SNRPB using two siRNAs. ***C***, HeLa cells were transfected with siRNA against SNRPB or with a non-targeting control siRNA (siCtrl). After 72 h, cells were treated with bortezomib (+) or solvent (-) followed by lysis and immunoblotting as indicated.

The reduction of the levels of ERES proteins was not due to a transcriptional effect because the mRNA levels of these proteins did not drop upon SNRPB depletion, but rather went up (e.g. Sar1A) (Figure 4B). The effect is also unlikely to be a consequence of the alteration of protein stability, because treatment of cells with bortezomib for 4 hours (an inhibitor of the proteasome) did not restore the levels of Sec24C, in contrast to β-catenin, which is known to be turned over by proteasomal degradation (Figure 4C).

### SNRPB mediates the splicing of Sec16A

Because SNRPB is a spliceosomal component, we hypothesized that downregulation of the levels of ERES proteins is due to defective splicing of their pre-mRNA. Among all affected proteins, we decided to focus on Sec16A, because it is a general regulator of ERES, that acts as an upstream regulator of all COPII components (Tang, 2017). To investigate a potential splicing defect of Sec16A (e.g. increased intron retention), we designed exon-based primers and amplified a region of the Sec16A, calnexin and actin transcripts. The primers were designed to cover exon 4-17 and would thus give rise to larger amplicons in case of impaired splicing due to intron retention (Figure 5A). Depletion of SNRPB resulted in a marked change in the pattern of bands for Sec16A and an appearance of a large band that matches the predicted size of an amplicon in the absence of any splicing (Figure 5A). The other bands are most likely partially spliced pre-mRNA. No effect was observed on actin or calnexin (Figure 5A), which is in line with the lack of an effect on SNRPB depletion on the protein levels of these genes (Figure 4A). We noticed that the SNRPB depleted condition showed some extra bands, which we consider to likely be splicing intermediates. Thus, it is likely that silencing SNRPB affects the protein levels of Sec16A due to impaired splicing. To further support these data, we reasoned that a splicing defect would result in a retention of the major fraction of Sec16A pre-mRNA in the nucleus. Therefore, we isolated RNA from nuclear and cytosolic fractions and determined Sec16A mRNA levels. The levels of Sec16A mRNA in SNRPB depleted cells were lower in the cytosolic fraction, and higher in the nuclear fraction (Figure 5B), which is in line with a splicing defect. Nuclear retention of pre-mRNA only occurs when it is associated with the spliceosome (Huang & Carmichael, 1996) which indicates that knockdown of SNRPB does not totally abolish formation of the spliceosome, but alters its function.

**Figure 5.**
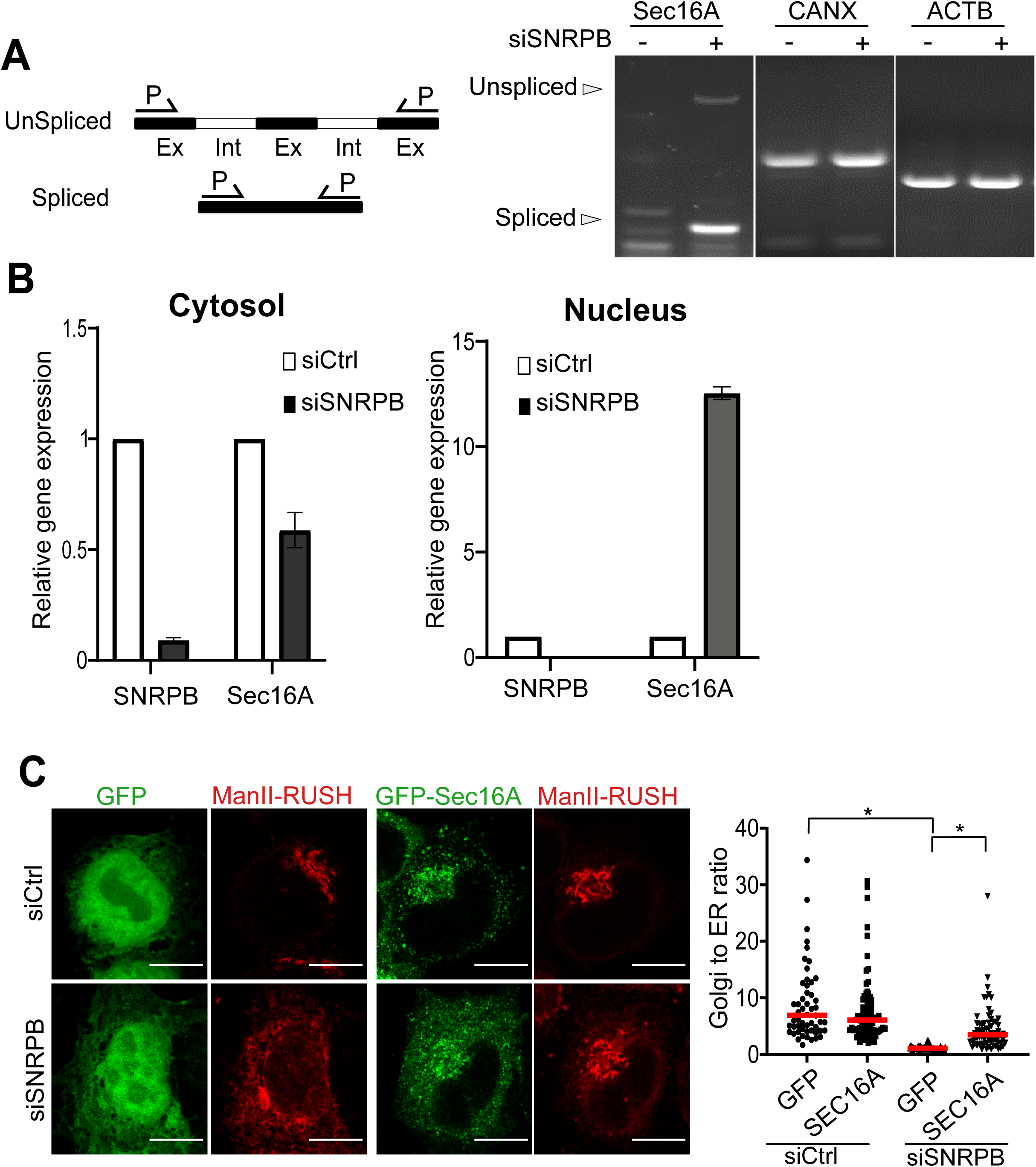
Depletion of SNRPB regulates Sec16A splicing. ***A***, RT-PCR using exon-based primers (spanning exons 4-17) as indicated in the schematic from cells depleted of SNRPB for 72 hours (EX= exons and Int= introns). RT-PCR was performed for the Sec16A, calnexin (CANX) and beta actin (ACTB). The two arrowheads indicate the positions of the spliced and non-spliced bands. ***B***, quantitative real time RT-PCR for SNRPB and Sec16A of cytosolic and nuclear fractions from cells transfected with non-targeting (siCtrl) or SNRPB siRNA or 72 h. ***C***, HeLa cells stably expressing mCherry tagged mannosidase-II RUSH construct (ManII) were transfected with siRNA against SNRPB (siSNRPB) or with a non-targeting control siRNA (siCtrl). After 48 h, cells were transfected with plasmids encoding GFP or GFP-tagged Sec16A. After 24 h, cells were treated with biotin for 20 min followed by fixation. The amount of mCherry signal in the Golgi area was normalized to the total cellular mCherry signal. The result of three independent experiments with 30 cells per experiment is displayed in the graph to the right. * indicates a p-value smaller than 0.05 obtained using one-way ANOVA.

To better link the effect of SNRPB on ER-export to its effect on Sec16A levels, we performed a rescue experiment using a GFP-tagged Sec16A construct. This construct is functional as shown earlier by us and others (Farhan *et al*., 2010; Tillmann *et al*, 2015). We overexpressed GFP-Sec16A in SNRPB depleted cells, and monitored ER-to-Golgi trafficking using the RUSH assay. As a control, we overexpressed GFP. Cells overexpressing GFP-Sec16A exhibited a partial restoration of ER-to-Golgi trafficking in SNRPB depleted cells (Figure 5C). These results indicate that the reduction in the levels of Sec16A in SNRPB silenced cells is the main factor that contributes to the defect in ER export.

### Global transcriptomic analysis reveals an effect of SNRPB on splicing of ERES regulators

To gain a systematic overview of the extent of the effects of SNRPB on the secretory pathway, we performed deep RNA sequencing of cells depleted of SNRPB. Samples were correctly assigned and exhibited strong silencing of SNRPB (Figure S3). First, we used Salmon and Deseq2 as an analysis pipeline for differential gene expression and the results are displayed as a volcano plot with genes related to secretory trafficking highlighted in green (Figure 6A). In line with our qPCR data on Sec16A and other components of the secretory pathway, we did not detect a change in mRNA levels of these genes. Next, we analyzed differential splicing using two pipelines: STAR/RMATS (Figure 6B, supplementary table 1) or the Whippet (Figure 6C, supplementary table 2). Both show that SNRPB knockdown results in massive cassette exon skipping as well as to a lesser extent intron retention, which is consistent with previous work (Saltzman *et al*, 2011). We plotted the changed exons in a volcano plot in Figure 6D, again highlighting exons in ER-Golgi trafficking genes in green. Overall, we identified 40 genes that exhibited affected transcripts. Expanding on these global analyses, manual screening of secretory genes revealed exon skipping and intron retention within a Sec16A region with low sequence coverage (Figure 6E), therefore escaping our global analysis. Overall, these data indicate that SNRPB regulates splicing of several secretory pathway genes and that Sec16A is one of the clients of this spliceosome that explains its effect on ERES and ER-to-Golgi trafficking. Among the affected transcripts were other components of the secretory pathway such as GOSR1, which we also found to exhibit reductions in protein levels similar to various ERES components (Figure S4). ERES proteins and GOSR1 were not affected by SNRPB2 depletion (Figure S4).

**Figure 6.**
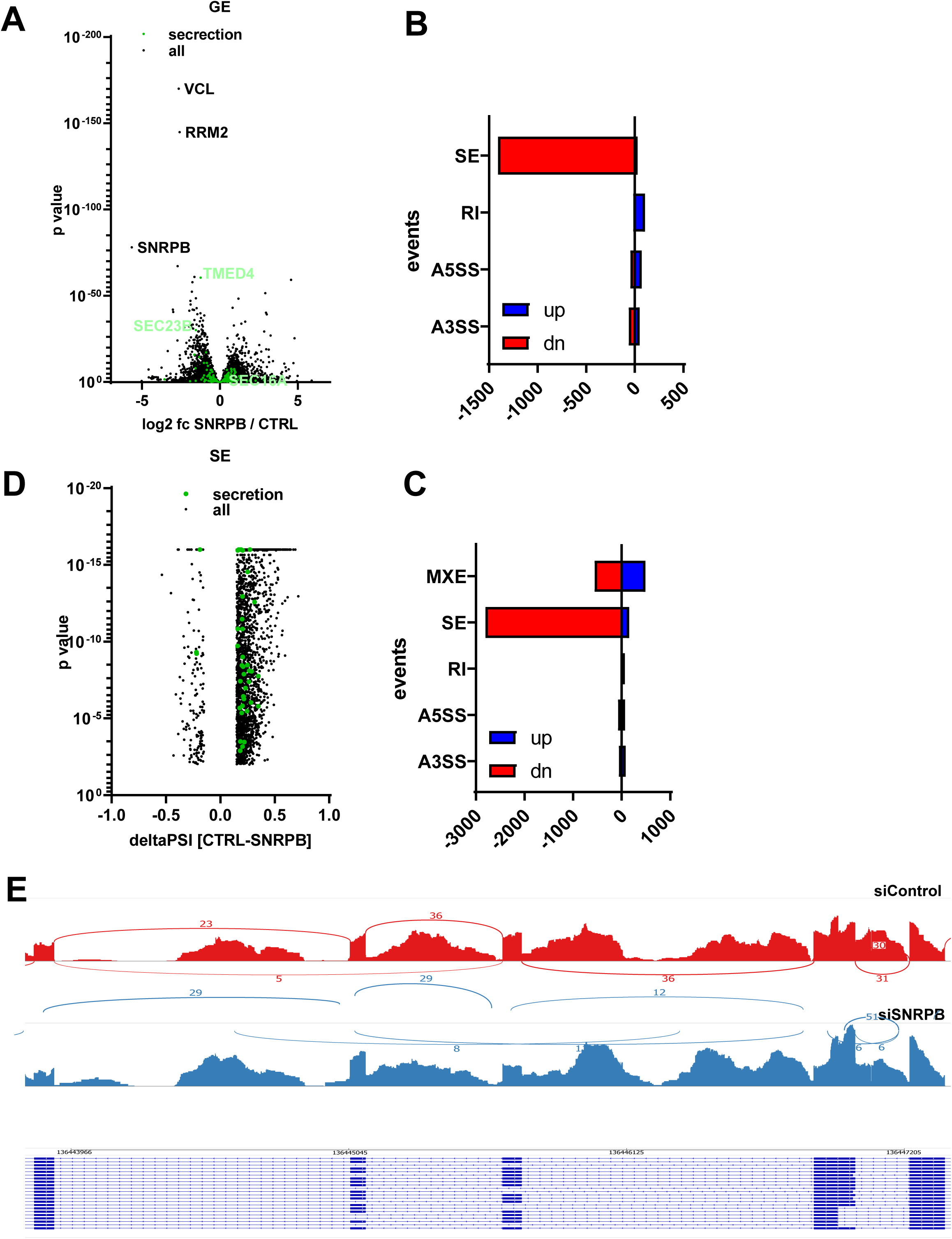
Systematic exploration of the effects of SNRPB depletion. ***A,*** Volcano blot highlighting SNRPB depletion induced changes in global gene expression (GE). Genes associated with the GO terms ‘COPII-coated vesicle budding’ and ‘regulation of regulated secretory pathway’ are highlighted in green. ***B, C,*** SNRPB depletion induced changes in alternative splicing derived by RMATS (***B***) or Whippet (***C***). Targets with a higher PSI upon knockdown are highlighted in blue, with a lower percent splice index (PSI) in red. MXE: mutually exclusive exons, SE: skipped exons, RI: retained introns, A3SS: alternative 3’splice-sites, A5SS: alternative 5’splice-sites. ***D,*** Volcano blot highlighting SNRPB depletion induced exon skipping. Exons in secretion associated genes are highlighted as in *A*. ***E,*** Sashimi plot highlighting SNRPB depletion induced changes in Sec16 splicing. siCTRL is shown in red, siSNRPB in blue. Pilled sequencing coverage is indicated on the y-axis. Junction reads are shown as lines.

We further determined whether there are common features in affected exons. We extracted 5 features from exons unregulated (n=108147), skipped (n=2804) or included (n=156) upon SNRPB knockdown. These features were upstream-intron, exon and downstream intron length, as well as upstream 3’ss and downstream 5’ss score. Exons skipped upon knockdown are slightly shorter and are surrounded by slightly shorter introns. Additionally, they are rather characterized by weak 5’ss than differences in 3’ss strength (Figure S5).

### SNRPB is regulated by cargo load in a manner dependent on ATF6

Our results so far suggest that SNRPB might regulate the ER-proteostasis network by modulating the function of ERES. We next wanted to gain a broader understanding of how SNRPB regulates ER-proteostasis, and whether SNRPB itself might be regulated by signaling pathways within the proteostasis network. The UPR is a transcriptional response that regulates ER export by inducing the expression of multiple ERES components. We hypothesized that spliceosomal subunit SNRPB might be involved in mediating the effects of the UPR on ERES. To induce UPR, we decided to use ER-overload, because we found in a previous study that the levels of Sec16A are elevated under conditions of ER-overload, in a manner dependent on the UPR (Farhan *et al*., 2008). To induce UPR by ER-overload, we overexpressed two constructs: GABA transporter 1 (GAT1) and GFP-b(5)tail. GAT1 is a polytopic transmembrane protein, which we showed previously to induce the UPR when overexpressed (Farhan *et al*., 2008). GFP-b(5)tail is a construct composed of the N-terminal cytosolic GFP moiety anchored to the ER membrane by the tail of cytochrome b5, which was shown previously to activate the ATF6 branch of the UPR (Maiuolo *et al*, 2011). We opted to induce UPR by overexpressing GAT1 and GFP-b(5)tail, instead of using chemical tools such as thapsigargin or tunicamycin, which do not accurately mimic the response to ER overload (Bergmann *et al*, 2018).

Overexpression of GAT1 as well as GFP-b(5)tail resulted in an induction of Sec16A levels, which was abrogated upon knockdown of SNRPB (Figure 7A). Furthermore, ER-overload resulted in an increase in the number of ERES, which was absent in SNRPB-depleted cells (Figure 7B). Similar results were obtained with GFP-b(5)tail overexpression, which increased the levels of Sec16A and the number of ERES in a manner dependent on SNRPB (Figure 7A&C).

**Figure 7.**
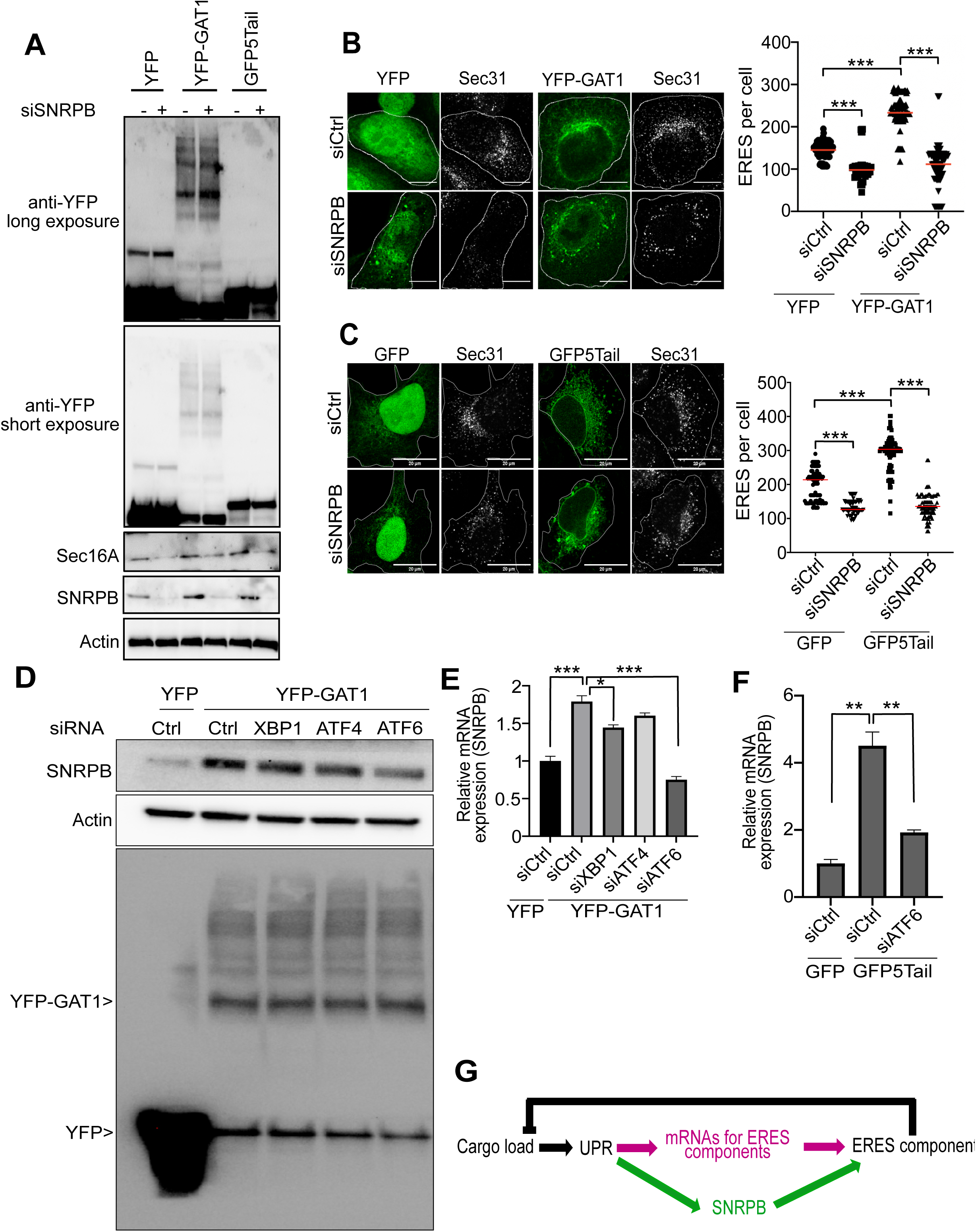
SNRPB is regulated by cargo load. ***A***, HeLa cells were transfected with siRNA against SNRPB (+) or with a non-targeting control siRNA (-). After 48 h, cells were transfected with a plasmid encoding YFP or YFP-tagged GAT1 or GFP-b(5)tail (GFP5Tail). After additional 24 h, cells were lysed and immunoblotted as indicated. ***B***, HeLa cells were transfected with siRNA against SNRPB (siSNRPB) or with a non-targeting control siRNA (siCtrl). After 48 h, cells were transfected with a plasmid encoding YFP or YFP-tagged GAT1. After additional 24 h, cells were fixed and immunostained against Sec31 to label ERES. The number of ERES was determined using ImageJ and the result of three independent experiments is displayed in the graph to the right. *** indicates a p-value smaller than 0.001 obtained using one-way ANOVA. ***C***, Same experimental setup as in panel B, except that cells were lysed transfected with GFP-b(5)tail. *** indicates a p-value smaller than 0.001 obtained using one-way ANOVA. ***D***, HeLa cells were transfected with siRNA against the indicated genes, or with a non-targeting control siRNA (siCtrl). After 48 h, cells were transfected with a plasmid encoding YFP or YFP-tagged GAT1. After additional 24 h, cells were lysed and immunoblotted as indicated. ***E***, Same as in panel D, except that we performed qPCR instead of a an immunoblot. Graph shows the levels of SNPRB mRNA normalized to siCtrl. *** indicates a p-value smaller than 0.001, and * indicates a p-values of smaller than 0.05, obtained using one-way ANOVA. ***F***, HeLa cells were transfected with siRNA against the indicated genes, or with a non-targeting control siRNA (siCtrl). After 48 h, cells were transfected with a plasmid encoding GFP or GFP-5tail. After additional 24 h, qPCR was performed to measure the levels of SNRPB mRNA. Graph shows the levels normalized to siCtrl with GFP expression. ** indicates a p-value smaller than 0.01 obtained using one-way ANOVA. ***G***, schematic depicting the role of SNRPB as apart pf a feed-forward loop in ER proteostasis.

We also noted that overloading the ER also resulted in a clear increase in SNRPB levels (Figure 7A), suggesting that SNRPB itself is regulated by the UPR. This would be in line with our hypothesis of SNRPB as a mediator or effector of the UPR. To test this hypothesis, we overexpressed GAT1 in cells depleted of XBP1, ATF4 and ATF6, which are main signaling branches of the UPR. GAT1 overexpression resulted in an increase in the protein levels of SNRPB in all conditions except in ATF6 knockdown cells (Figure 7D). The induction of SNRPB was also visible at the mRNA level, which is in agreement with the UPR being a transcriptional response. Again, the induction of SNRPB by ER-overload was abrogated in ATF6 depleted cells (Figure 7E). Silencing of XBP1 or ATF4 had only a very weak effect indicating that ATF6 is the major pathway that is used to induce SNRPB. Finally, we found that GFP-b(5)tail overexpression induces the mRNA levels of SNRPB, which was abrogated by silencing ATF6 (Figure 7F).

Overall, these results indicate that SNRPB is an effector of the UPR, while at the same time being a facilitator of the effects of the UPR on ER proteostasis (Figure 7G).

### Loss of SNRPB induces osteogenesis defects in vitro that are rescued by Sec16A overexpression

Loss of function of SNRPB underlies the rare genetic disease CCMS, which is characterized by severe bone deformations, microcephaly, rib defects and micrognathia (Bacrot *et al*., 2015; Leroy *et al*, 1981; McNicholl *et al*, 1970). While we are fully aware that bone development is complex process that is regulated by multiple factors, we asked whether the effect of SNRPB on ER-proteostasis might contribute partially to the pathophysiology of CCMS. This conjecture is strengthened by the notion that alterations of multiple proteins of the ER-export machinery were associated with bone defects (Boyadjiev *et al*., 2006; El-Gazzar *et al*., 2023; Garbes *et al*., 2015). We therefore tested whether loss of SNRPB affects osteogenesis using an in vitro assay where we differentiated mesenchymal cells to osteoblasts. Osteogenesis can be visualized using alizarin red staining, which detects calcification. Silencing SNRPB with two siRNAs resulted in a notable reduction in osteogenesis, as indicated by decreased alizarin red staining intensity (Figure 8A-C).

**Figure 8.**
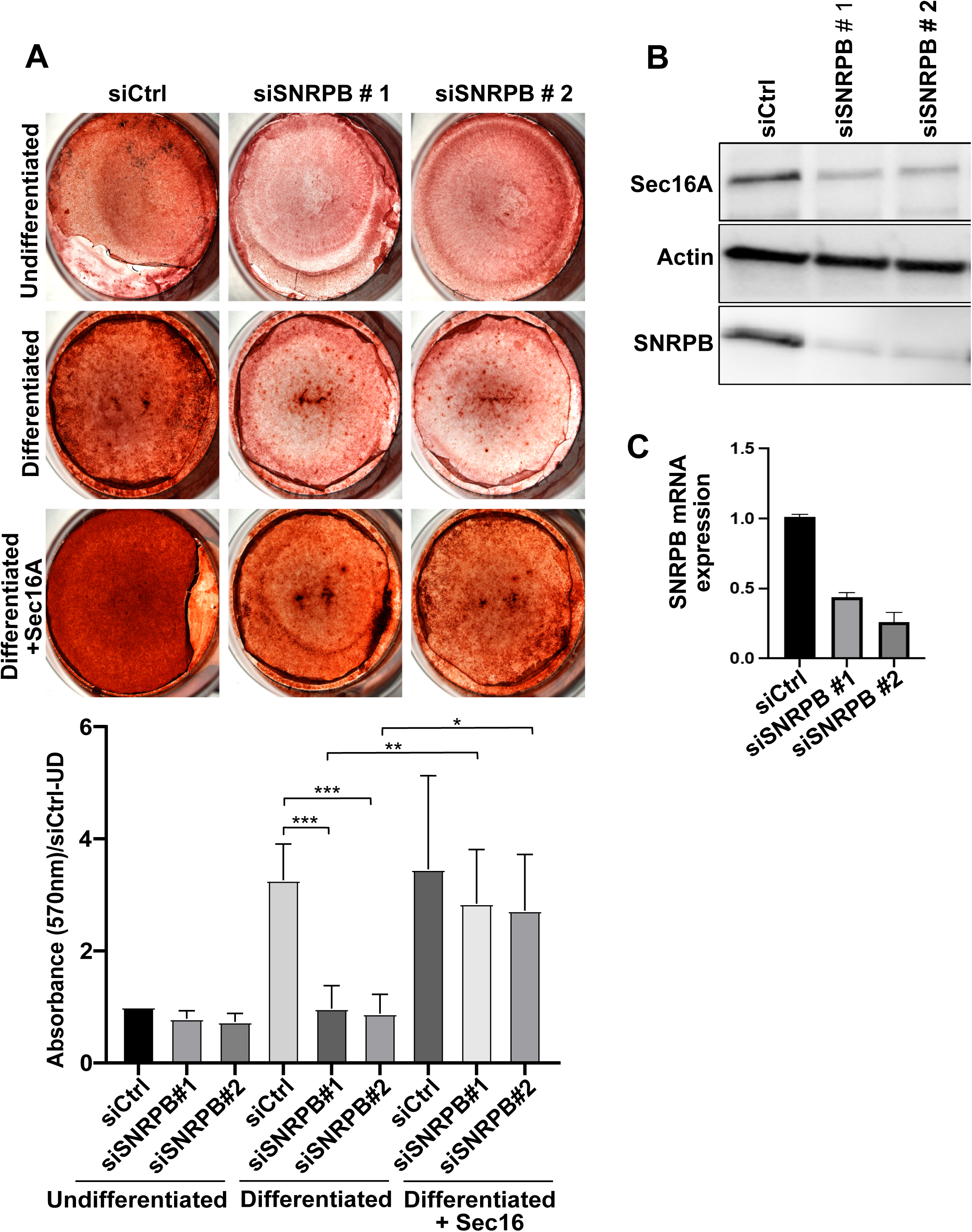
Loss of SNRPB results in osteogenesis defect in vitro, which is rescued by Sec16A. ***A***, Mesenchymal cells were transfected with non-targeting siRNA (Ctrl) or with two siRNAs against SNRPB. Cells were subjected osteogenic differentiation protocol for 9 days followed by alizarin red staining. ***B&C***, demonstration of knockdown efficiency of the SNRPB siRNAs in mesenchymal cells using immunoblotting (B) and qPCR (C).

To link this effect to ERES function, we asked whether overexpression of Sec16A might rescue the effect of SNRPB depletion. We chose Sec16A because it plays a key role in the biogenesis and maintenance of ERES, upstream of COPII subunits. In cells with reduced levels of COPII components, overexpression of Sec16A would potentiate the recruitment of the available COPII subunits to ERES to drive ER export. In our previous work, we showed that signaling to Sec16A is sufficient to drive biogenesis of new ERES without the need to regulate the levels of any COPII component (Tillmann *et al*., 2015). As shown in Figure 8A, overexpression of Sec16A almost completely rescued the osteogenesis defect in SNRPB-depleted cells. These results indicate that the bone defects observed under loss of SNRPB is, at least partially, linked to a defect in the ERES function.

### Inducible Snrpb Deletion Results in Chondrogenesis and Osteogenesis Defects

We previously showed that embryos with heterozygous loss of function mutation in Snrpb arrested at embryonic day (E) 8.5 (Alam *et al*., 2022), indicating an early requirement for this gene during gastrulation. We also showed bone deformities in the craniofacial region of neural-crest cell Snrpb mutants. To examine the requirement for Snrpb in bone development in the appendages, we used tamoxifen to induce deletion of Snrpb at E8.5. Mice carrying LoxP sequences flanking exon 2 and 3 of *Snrpb* (Snrpb^loxP/+)^ (23) and ER-Cre^tg/tg^ (Ventura *et al*, 2007) were crossed to produce Snrpb^loxP/+;^ *ER-Cret^g/tg^* males for mating with wild type CD1/CD57 females. Pregnant females were injected with a single dose of tamoxifen at E8.5 to induce deletion of exons 2 and 3 of *Snrpb*. The resulting litters were collected at E14.5 (n= 18 controls and 14 mutants) and E17.5 (n= 15 controls and 12 mutants) and stained with Alcian blue and Alizarin red to visualize the cartilage and bones, respectively. Below, we describe severe malformations in both limbs and girdles (scapula and pelvis) of *Snrpb^LoxP/+;^ER-Cre^Tg/+^*embryos at E14.5 and E17.5 (Figure 9A-F). The scapula was smaller and narrower in most mutant embryos (n=19/26) and showed dysmorphologies including a missing blade (n=7/26), bifurcated blade (n=7/26), and fused shoulder joints (n=10/26). In a small subset of E17.5 embryos, a curvy clavicle (n=4/12) was also found. In the pelvic, the iliac bone was thinner (n=18/26) or missing (n= 9/26) in Snrpb mutant embryos. Defects were also found in the stylopod. The humerus of Snrpb mutants was narrower in E14.5 embryos (n=3/14) than those of controls and few were identified with missing the deltoid tuberosity (n=2/14). At E17.5, the deltoid tuberosity was misshapen or smaller in six of twelve mutant embryos analyzed. In addition, the olecranon process was malformed in mutants of both stages (n=4/26). The femur of Snrpb mutants was also thinner and had an abnormal narrowing at the middle (n= 8/26). The zeugopod of mutants was often bent, although this was more evident in the ulna (n=19/26) when compared to the radius (n=17/26). Additionally, fewer embryos had bending in the tibia and fibula (n=6/26). Defects, in the autopod included fused 2nd and 3rd metacarpal (n=1/26), bilaterally, and a missing 5th digit (n=1/26). However, these malformations were rare, with each found in one Snrpb mutant embryo. Moreover, clinodactyly, or bent 5th phalanges was found in both controls (n=4/33) and mutant (n =4/26) embryos. In addition to these dysmorphologies, at E17.5, Alizarin red staining was missing in the ulna and radius of six mutant and four control embryos, or split into two domains in the ulna of one mutant embryo, indicating abnormal osteogenesis. Finally, the humerus, radius, ulna, and femur of mutants (n=12) were shorter when compared to those of controls (n=15) (unpaired, two-tailed t-test, P<0.05) (Figure 9G). Hence, reducing levels of Snrpb after a single injection of tamoxifen at E8.5, leads to malformations in appendage of the majority of *Snrpb^LoxP/+;^ER-Cre^Tg/+^* embryos (n=25/26), except for one E14.5 mutant embryo. Furthermore, in most E14.5 (n=10/14) and all E17.5 (n=12) mutant embryos these malformations were bilateral, though the severity differed between the left and right side. Altogether our data indicate that normal level of Snrpb is required for growth and patterning of the proximal and distal elements of the mouse appendage, and for both chondrogenesis and osteogenesis.

**Figure 9.**
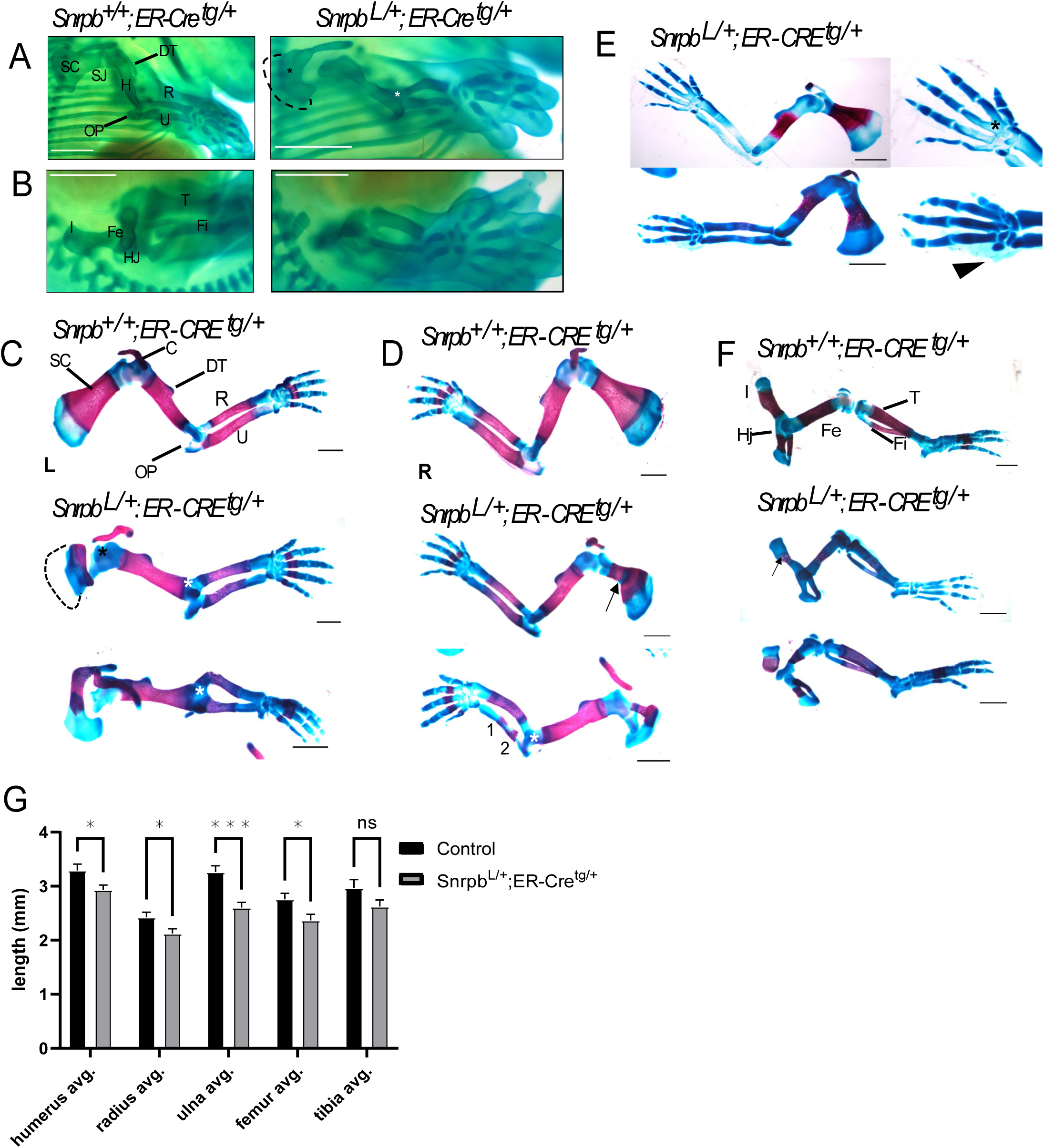
Tamoxifen-induced *Snrpb* deletion before E8.5 causes chondrogenesis and osteogenesis defects in appendages. Representative images of Alcian Blue-strained E14.5 control (*Snrpb^+/+^; ER-Cre^tg/+^*) and mutant (*Snrpb^L/+^; ER-Cre^tg/+^*) embryos showing chondrogenesis defects in (A) forelimbs and (B) hindlimbs. ***A,*** Dorsal-lateral view of right forelimbs showing missing shoulder blade (dashed line), fused shoulder joint (SJ, black asterisk), narrow humerus (H), absent deltoid tuberosity (DT), fused elbow joint (EJ, white asterisk), malformed olecranon process (OP), and bent radius (R) in the mutant. ***B,*** Dorsal-lateral view of right hindlimbs showing missing iliac bone (I), narrow femur (Fe), bent tibia (T) and fibula (Fi) in the mutant. Representative images of Alcian Blue-and Alizarin Red-stained E17.5 control and mutant embryos showing chondrogenesis and osteogenesis defects in the left (C) and right (D-E) forelimbs and the right hindlimbs (F). ***C,*** Ventral-lateral view of left forelimbs showing missing scapula blade (dashed line), curved clavicle (C), fused shoulder joint (black asterisk), misshapen deltoid tuberosity, fused elbow joint (white asterisk), malformed olecranon process, bent radius and ulna (U) in mutants. ***D,*** Ventral-lateral view of right forelimbs showing smaller and narrower scapula (SC, arrow), bifurcated blade (double curved arrow), curved clavicle (C), misshapen deltoid tuberosity, fused elbow joint (white asterisk), malformed olecranon process, bent radius, and abnormal ulna ossification (two domains labelled 1 and 2) in mutants. ***E,*** Ventral-lateral view of right forelimbs showing smaller and narrower scapula, missing Alizarin red staining in the radius and ulna (top), fused 2^nd^ and 3^rd^ metacarpals (black asterisk), clinodactyly (arrow), and missing 5^th^ digit (arrowhead) in mutants. ***F,*** Dorsal-lateral view of right hindlimbs showing thin (top, arrow) and missing iliac bone (bottom), narrow femur (bottom), and bent tibia (bottom) in mutants. Scale bar=1mm. ***G,*** Length of the humerus, radius, ulna, and femur is reduced in *Snrpb^L/+^; ER-Cre^tg/+^* mutants (unpaired, two-tailed *t*-test). \**P*<0.05; \*\*\**P*<0.0001. Error bars indicate SEM.

## Discussion

Research over the past few decades has expanded our understanding of the role of the UPR in regulating cellular proteostasis under physiological and pathological conditions such as cancer and neurodegenerative disorders (Hetz & Saxena, 2017; Urra *et al*, 2016). The UPR is a transcriptional response and so far, it has been assumed that no regulatory step exists between the UPR-mediated induction of mRNAs and their translation into proteins. This was based on the assumption that the splicing machinery is not a rate-limiting factor in this process. Our current work indicates that this assumption is not correct and expands our understanding of the regulation of ER proteostasis. We show that the spliceosomal component SNRPB is required for the adaptation of the ER to higher loads of secretory cargo where it acts as part of a feed-forward loop that signals from UPR to ERES in an ATF6-dependent manner. In the absence of SNRPB, cargo load failed to induce an increase in the levels of the ER-export machinery. We hypothesize that the induction of several hundred mRNAs by the UPR exceeds the capacity of the pre-existing pool of SNRPB, thus necessitating the induction of this spliceosomal component by ATF6.

Mechanistically, the effect of SNRPB on ER proteostasis was mediated through splicing of components of the membrane trafficking machinery such as regulators of biogenesis and trafficking at ERES. Consequently, silencing of SNRPB reduced the number of ERES and delayed trafficking from the ER to the Golgi. Although several ERES components were affected, we were able to rescue the ERES defect by exogenous expression of Sec16A, indicating that this might represent the dominant component behind the effect of SNRPB on ER proteostasis. In the absence of SNRPB, cells failed to upregulate the levels of Sec16A in response to ER overload. Thus, we propose SNRPB as a new checkpoint in the ER-proteostasis network. We noted that the regulatory circuit has features of a coherent feed-forward loop (CFFL) (Figure 7G). The biological function of many CFFLs is to act as persistence detectors to ensure that only a persistent stimulus exerts a full biological response (Lim *et al*, 2013). A feature of CFFLs is that one of the two forward loops is slower than the other. In our case, the loop that includes SNRPB (green in Figure 7G) contributes in a slower manner towards the output than the loop where the UPR stimulates the transcription of ERES components (purple in Figure 7G). We hypothesize that the initial UPR response is handled by the pre-existing pool of SNRPB. However, a persistent response creates more clients for SNRPB and therefore necessitates higher levels of this spliceosomal component. This is supported by the observation that SNRPB knockdown cells fail to induce the levels of Sec16A under condition of cargo overload. Thus, our data identify a role for core splicing in the regulation of ER-proteostasis.

Mutations in SNRPB were shown to be linked to a rare genetic disease called Cerebro-costo-mandibular syndrome (CCMS). Skeletal abnormalities are a hallmark of CCMS, but there was no obvious link between mutations in a spliceosomal component and bone abnormalities. Our results provide mechanistic insights into the pathophysiology of the disease. Based on our observations, we propose that the mechanism of bone deformations is due to a perturbation of ER homeostasis and the trafficking of collagen-I. Previous work has demonstrated several examples where mutations in genes regulating ER homeostasis result in bone deformation as has been shown for Sec23A, Sec24D, and ATF6 (Boyadjiev *et al*., 2006; Garbes *et al*., 2015; Guo *et al*, 2016). Further support for this notion comes from our experiment in patient-derived fibroblasts where we could increase the number of ERES by expressing wild type SNRPB. We are aware that it is difficult to deduce clinically relevant effects based on cell culture experiments, but this result opens the possibility that a defect in ER export might be associated with the bone developmental defects in CCMS patients.

Altogether, our work establishes a regulatory role for the core spliceosomal component SNRPB in ER-proteostasis and provides insights into the pathophysiology of Cerebro-costo-mandibular syndrome.

## Materials and Methods

### Cell culture and transfection

Hela cells, U2OS and Hela-Rush cells (stably expressing ManII-GFP RUSH plasmid) were cultured at 37oC, 5% CO2 and normal humidity. For confocal microscopy, cells were seeded at a density of 2×10^5^ cells per well in a 6-well plate. All siRNAs used in this study are listed in Supplementary Table S2. Depletions were performed with siRNA from Thermofisher Scientific (silencer select) and independently validated with siRNA from Dharmacon (Dharmacon siGenome-SMARTpool). The non-targeting siRNA from Thermofisher Scientific (silencer select) was used as the siRNA control. Briefly, 10 nM siRNAs were mixed with 6 µl of HiPerFect transfection reagent (Qiagen, #301707) in 100 µl of serum free DMEM and added to freshly plated cells drop by drop. Plasmids were transiently overexpressed using TransIT-LT1 (Mirus, MIR2306) according to the instructions of the manufacturer. In short, 750ng of DNA was transfected per well, in a 6-wells plate for 1 day using 3µl of Mirus per 1µg of DNA. For collagen-1 experiments, 50ng/ml of ascorbic acid was treated for 6 hours.

### Immunofluorence staining and confocal microscopy

Cells grown on glass coverslips were fix in 4% PFA for 6 min, washed with PBS, incubated in 50 mM NH4Cl for 8 min at room temperature and then washed with PBS. Next, the cells were incubated in permeabilization buffer (0.25% Triton X-100 in PBS) for 8 min at room temperature, washed with washing buffer (0.05% tween-20) and then incubated with blocking buffer (5% BSA in washing buffer) for 1 h at room temperature. Cells were incubated with primary antibodies in 1% BSA (in washing buffer) for 1 h at room temperature. After this incubation, the cells were washed with washing buffer and incubated for 1 h with secondary antibodies in 1% BSA. The coverslips were briefly rinsed with Milli-Q water and embedded in polyvinyl alcohol mounting medium (Sigma Aldrich, catalog # 10981). Fluorescence images were acquired on a LSM 700 confocal microscope using a PlanAphochromat 63x/1.40 Oil Ph3 M27 objective. For ERES experiments, images were taken, focusing at nuclear ring and adjust the threshold level, when only puncta siganal is detected. The number of ERES were counted per cell. The list of all antibodies including dilutions is provided under Key Resources below.

### Immunoblotting

Briefly, cells were lysed in RIPA buffer supplemented with cocktail of protease and phosphatase inhibitors (Theremofisher; A32961). The blots were incubated with primary antibodies overnight in cold room, washed with PBST and subsequently incubated with HRP secondary antibodies for 1 hour at room temperature.

### RNA isolation and qPCR

To measure mRNA, cells were transfected with siRNAs/plasmid as described earlier and RNA was extracted using GenElute™ Mammalian Total RNA Miniprep Kit (Sigma Aldrich, # RTN70). RNA was reverse transcribe using High-Capacity cDNA Reverse Transcription Kit (Thermo Fisher #4368814). Real-time PCR was performed with 9ng of cDNA per reaction using Quantitect primers (Qiagen).

### Nuclear Fractionation

Cells were incubated with hypotonic solution containing Hepes (pH 7.5, 10mM), MgCl2 (2mM), KCl (25mM) and cocktail of protease/phosphatase inhibitors for 1 hours. Cell were broken using Pellet Pestle Motor Kontes (Thomas Scientific) and Disposable Polypropylene, RNase-Free Pellet Pestles (Thomas Scientific) for 15 second with a break every 5 second. The cells were checked under light microscope for nuclear integrity. After homogenization, 2M sucrose solution was added at concentration of 125µl/ml and then centrifuged at 600xg for 10 minutes in swinging bucket router. Pellet contains nuclei and are washed twice with hypotonic solution containing sucrose. Pellet and supernatant are used for RNA solution same as above. The whole protocol is performed at 4°C.

### RNA-Seq analysis

Libraries were sequenced on an Illumina HiSeq3000 flow cell, using 2-lanes. This yielded ∼ 35-40 million paired-end 150 nt reads for biological triplicates and technical duplicates of siCTRL and siSNRPB samples. These were analyzed for gene expression and alternative splicing using independent pipelines. For differential gene expression reads were aligned to the human genome using salmon (version 1.5.1) (Patro *et al*, 2017). Salmon quantifications of gene expression were imported into RStudio using tximport and analyzed for differential gene expression using Deseq2 (version 1.12.3) (Love *et al*, 2014). Genes associated with the GO terms ‘COPII-coated vesicle budding’ and ‘regulation of regulated secretory pathway’ were downloaded from Panther GO and merged with the Deseq2 output using standard Python code. For alternative splicing two independent splicing pipelines were used. First, reads were aligned to the human hg38 genome, using STAR (version 2.7.9a) (Dobin *et al*, 2013), yielding ∼ 85 % uniquely aligned reads. Alternative splicing changes were calculated using rMATS (version v3.1.0) (Shen *et al*, 2014). To obtain only high confidence targets, only targets with a p-value < 0.01 and a deltaPSI > 0.15 were considered alternatively spliced. Additionally, to filter out splicing events in weakly expressed genes or gene regions with low expression, events with less than 100 combined junction reads in all samples were excluded. Second, alternative splicing changes were quantified – independent rom the STAR/RMATS pipeline – using Whippet (v0.11.1) (Sterne-Weiler *et al*, 2018). High confidence targets were filtered on Whippet-delta derived Probability > 0.95 and a deltaPSI > 0.15. To investigate exon features, exon and intron-length were calculated using standard Python code. Splice site coordinates were extracted from the rMATS output table in bed format (-20 to +3 for 3’-ss; -3 to +6 for 5’- ss) and splice site sequences were extracted using bedtools. Sequences were input to MaxEntScan (Yeo & Burge, 2004), using the Maximum Entropy Model to score.

### Osteogenic differentiation

MC3T3-E1 cells were cultured in MEM-α media, supplemented with 10% FBS under normal humidity and 5% CO2. For the osteogenic differentiation of MC3T3, cells were cultured in osteogenic media (DMEM media supplemented with 10% FBS, 50µg/ml of ascorbic acid and 1mM β-glycerophosphate for 18 days. The media were changed every 3 days. For Alizrin red staining, cells were fixed with PFA (4%) and gently washed with dH2O. The cells were then incubated with 40mM (pH 4.3) alizarin red for 2 hours at room temperature and then washed with dH2O with gentle shaking. For the quantification of the alizarin red staining, the wells were incubated with 35mg/ml of Cetylpyridinium Chloride (CPC) for 2 hours at 37°C until fully dissolved. The solution was collected, cleared by brief centrifugation and diluted in CPC and then measured at 570 nm.

For the knockdown experiments, plates were coated with siRNA using Lipofectamine 2000 as transfection reagent. After 24 h, GFP-Sec16A plasmid was transfected and cells were allowed to express the protein for 24h. Afterwards, media were changed to differentiation media and cells were allowed to differentiate for 18 days followed by Alizarin red staining as indicated above.

### Mouse lines

All procedures and experiments were performed according to the guidelines of the Canadian Council on Animal Care and approved by the animal Care Committee of the Montreal Children’s Hospital. Wnt1-Cre2 (Lewis *et al*, 2013) mice on the C57BL/6J genetic background were purchased from The Jackson Laboratory (strain#: 022501). The development of a conditional knock-out Snrpb allele with CRISPR/Cas9-mediated homology directed repair (HDR) strategy was previously described (Alam *et al*., 2022).

### Generation of neural crest cell-specific Snrpb^+/-^ mutant embryos (Snrpb^ncc+/-^)

To generate embryos with neural crest-specific Snrpb heterozygosity, Wnt1-Cre2^tg/+^ animals were mated with Snrpb^loxp/+^ mice. Embryos obtained from these mating were Snrpb heterozygous mutant in the neural crest cells and their derivatives, while all other cells were Snrpb wild type.

### Collection and genotyping of embryos

The day of plug was considered embryonic day 0.5 (E0.5). Embryos were collected at E9.5. On the day of dissection embryos were removed from their extraembryonic membranes and assessed for the presence of a heartbeat. The somite number was counted under light microscope (Leica MZ6 Infinity1 stereomicroscope) at the time of dissection. Embryos were fixed in 4% paraformaldehyde in PBS at 4°C overnight, washed and kept in PBS at 4°C until use. Yolk sacs were used for genomic DNA extraction and genotyping (Hou *et al*, 2017). Genotyping to identify Snrpb wild-type and conditional allele (with loxP sequences) was previously described (Alam *et al*., 2022). The primers used for the genotyping were: forward-5’ CCCGAGACAGACACAACATAAG 3’, reverse-5’ GCTTTGAAGGTCCCGATGAA 3’.

### Tamoxifen Treatment

In the morning of E8.5, pregnant females were injected with a single dose Tamoxifen (Sigma, T5648), 2 mg per 20 g mice, intra-peritoneally. Tamoxifen was dissolved at a concentration of 10mg/ml in sterile corn oil.

### Cartilage and skeletal preparation of embryos

To investigate cartilage formation, embryos were stained with Alcian Blue. For skeletal staining, the skin was removed from freshly dissected E17.5 embryos and neonatal pups and stained as described by (Wallin *et al*, 1994). Cartilage and skeletal preparations were imaged and analyzed using Lecia stereomicroscope (Lecia MZ6). Measurements were taken using Fiji ImageJ and statistical analysis was carried out on Graphpad, Prism (https://www.graphpad.com/scientific-software/prism/).

### Preparation of embryos for embedding and histology

For cryo-embedding, fixed embryos were first cryoprotected in 30% sucrose overnight, embedded in cryomatrix and sectioned at 10mm thickness for immunofluorescence.

### Immunofluorescence of tissue section

After three 10 min washes with phosphate-buffered saline (PBS), slides were microwaved for 1 min at 100% power and 3 min at 20% power in 10 mM Na Citrate (pH6) for antigen-retrieval. Once returned to room temperature, slides were blocked at 4 °C overnight with blocking buffer, 10% horse serum in PBS containing 0.25% Triton X-100. Slides were then incubated with Anti-Type I Collagen (1:100; SouthernBiotech; Cat. # 310-01) primary antibody, diluted in blocking buffer, at 4 °C overnight. After washes with PBS, slides were incubated with fluorescein-conjugated secondary antibody Rabbit anti-Goat IgG (H+L) Cross-Adsorbed

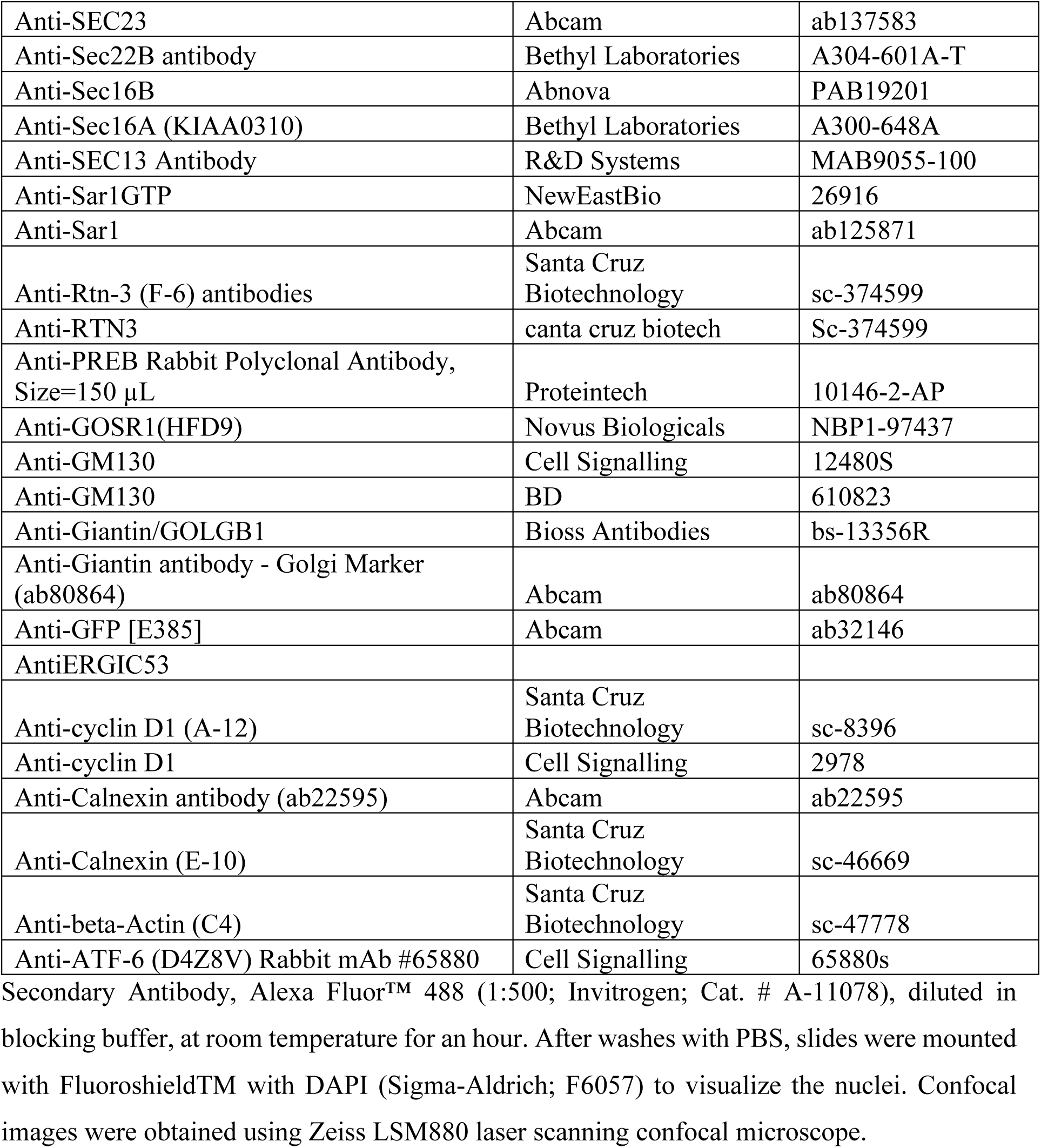

|  |  |  |
| --- | --- | --- |
| Anti-SEC23 | Abcam | ab137583 |
| Anti-Sec22B antibody | Bethyl Laboratories | A304-601A-T |
| Anti-Sec16B | Abnova | PAB19201 |
| Anti-Sec16A (KIAA0310) | Bethyl Laboratories | A300-648A |
| Anti-SEC13 Antibody | R&D Systems | MAB9055-100 |
| Anti-Sar1GTP | NewEastBio | 26916 |
| Anti-Sar1 | Abcam | ab125871 |
| Anti-Rtn-3 (F-6) antibodies | Santa Cruz Biotechnology | sc-374599 |
| Anti-RTN3 | canta cruz biotech | Sc-374599 |
| Anti-PREB Rabbit Polyclonal Antibody, Size=150 µL | Proteintech | 10146-2-AP |
| Anti-GOSR1(HFD9) | Novus Biologicals | NBP1-97437 |
| Anti-GM130 | Cell Signalling | 12480S |
| Anti-GM130 | BD | 610823 |
| Anti-Giantin/GOLGB1 | Bioss Antibodies | bs-13356R |
| Anti-Giantin antibody - Golgi Marker (ab80864) | Abcam | ab80864 |
| Anti-GFP [E385] | Abcam | ab32146 |
| AntiERGIC53 |  |  |
| Anti-cyclin D1 (A-12) | Santa Cruz Biotechnology | sc-8396 |
| Anti-cyclin D1 | Cell Signalling | 2978 |
| Anti-Calnexin antibody (ab22595) | Abcam | ab22595 |
| Anti-Calnexin (E-10) | Santa Cruz Biotechnology | sc-46669 |
| Anti-beta-Actin (C4) | Santa Cruz Biotechnology | sc-47778 |
| Anti-ATF-6 (D4Z8V) Rabbit mAb #65880 | Cell Signalling | 65880s |

### Key resources used

#### Antibodies

#### Chemicals

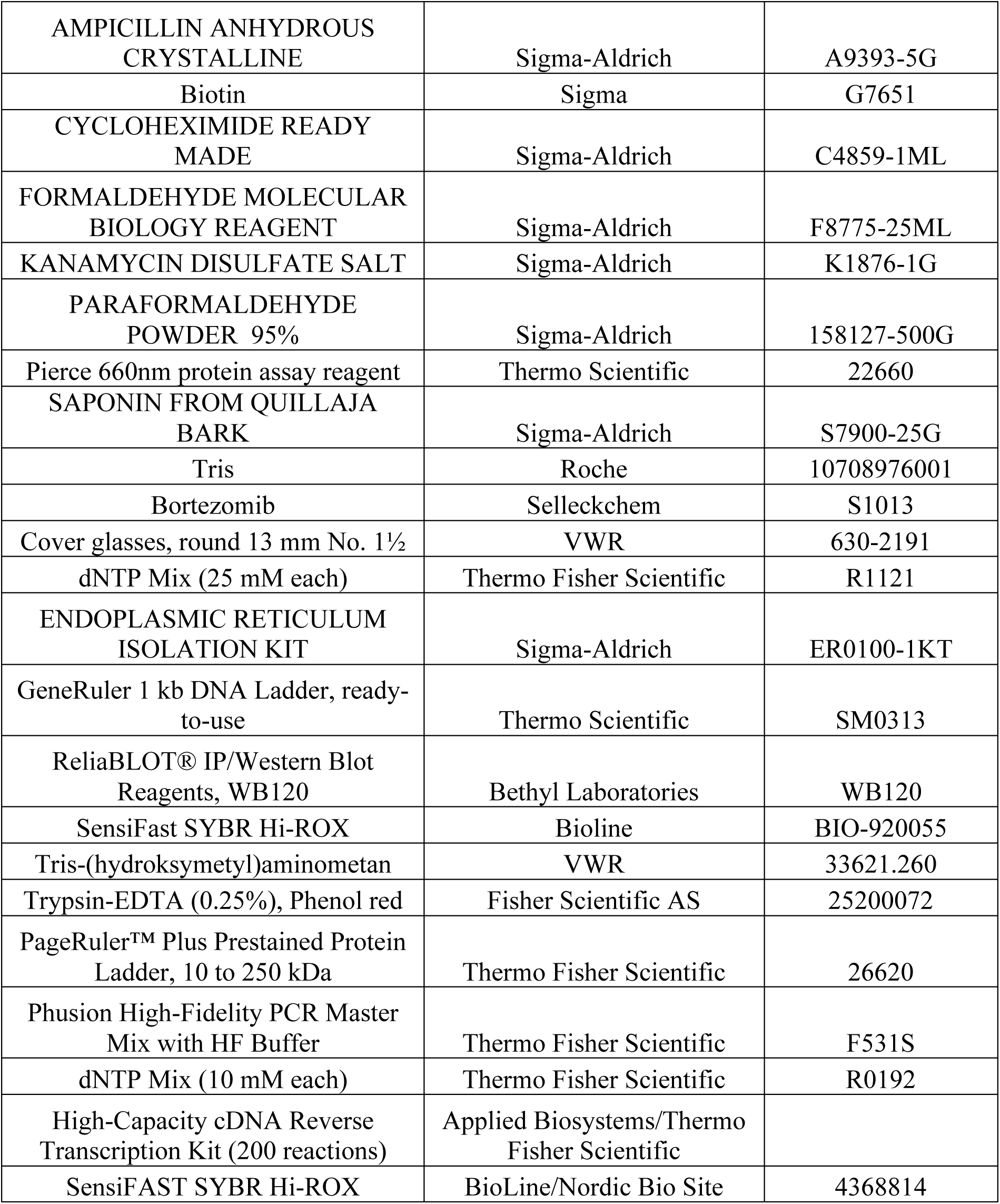

|  |  |  |
| --- | --- | --- |
| AMPICILLIN ANHYDROUS CRYSTALLINE | Sigma-Aldrich | A9393-5G |
| Biotin | Sigma | G7651 |
| CYCLOHEXIMIDE READY MADE | Sigma-Aldrich | C4859-1ML |
| FORMALDEHYDE MOLECULAR BIOLOGY REAGENT | Sigma-Aldrich | F8775-25ML |
| KANAMYCIN DISULFATE SALT | Sigma-Aldrich | K1876-1G |
| PARAFORMALDEHYDE POWDER 95% | Sigma-Aldrich | 158127-500G |
| Pierce 660nm protein assay reagent | Thermo Scientific | 22660 |
| SAPONIN FROM QUILLAJA BARK | Sigma-Aldrich | S7900-25G |
| Tris | Roche | 10708976001 |
| Bortezomib | Selleckchem | S1013 |
| Cover glasses, round 13 mm No. 1½ | VWR | 630-2191 |
| dNTP Mix (25 mM each) | Thermo Fisher Scientific | R1121 |
| ENDOPLASMIC RETICULUM ISOLATION KIT | Sigma-Aldrich | ER0100-1KT |
| GeneRuler 1 kb DNA Ladder, ready-to-use | Thermo Scientific | SM0313 |
| ReliaBLOT® IP/Western Blot Reagents, WB120 | Bethyl Laboratories | WB120 |
| SensiFast SYBR Hi-ROX | Bioline | BIO-920055 |
| Tris-(hydroksymetyl)aminometan | VWR | 33621.260 |
| Trypsin-EDTA (0.25%), Phenol red | Fisher Scientific AS | 25200072 |
| PageRuler™ Plus Prestained Protein Ladder, 10 to 250 kDa | Thermo Fisher Scientific | 26620 |
| Phusion High-Fidelity PCR Master Mix with HF Buffer | Thermo Fisher Scientific | F531S |
| dNTP Mix (10 mM each) | Thermo Fisher Scientific | R0192 |
| High-Capacity cDNA Reverse Transcription Kit (200 reactions) | Applied Biosystems/Thermo Fisher Scientific |  |
| SensiFAST SYBR Hi-ROX | BioLine/Nordic Bio Site | 4368814 |

#### qPCR primers

|  |  |  |
| --- | --- | --- |
| Hs_ACTB_2_SG QuantiTect Primer Assay | Qiagen | QT01680476 |
|  | Qiagen | QT01680476 |
|  | Qiagen | QT01680476 |
| Hs_CANX_1_SG QuantiTect Primer Assay | Qiagen | QT00092995 |
| Hs_GOSR1_1_SG QuantiTect Primer Assay | Qiagen | QT00069447 |
|  | Qiagen | QT00096404 |
| Hs_LOC442272_1_SG QuantiTect Primer Assay | Qiagen | QT00277718 |
| Hs_PREB_1_SG QuantiTect Primer Assay | Qiagen | QT00209958 |
| Hs_SAR1B_1_SG QuantiTect Primer Assay | Qiagen | QT00039865 |
| Hs_SEC16A_1_SG QuantiTect Primer Assay | Qiagen | QT00092778 |
| Hs_SEC23A_1_SG QuantiTect Primer Assay | Qiagen | QT00048468 |
| Hs_SEC24A_1_SG QuantiTect Primer Assay | Qiagen | QT00215859 |
| Hs_SEC24B_1_SG QuantiTect Primer Assay | Qiagen | QT00048664 |
| Hs_SEC24C_1_SG QuantiTect Primer Assay | Qiagen | QT00014336 |
| Hs_SEC24D_1_SG QuantiTect Primer Assay | Qiagen | QT00042959 |
| Hs_SNRPA1_1_SG QuantiTect Primer Assay | Qiagen | QT00094003 |
| Hs_SNRPB_1_SG QuantiTect Primer Assay | Qiagen | QT00014686 |
| Hs_SNRPB2_1_SG QuantiTect Primer Assay | qiagen | QT00035994 |
| Hs_SNRPC_1_SG QuantiTect Primer Assay | Qiagen | QT00000707 |
| Hs_SNRPD1_1_SG QuantiTect Primer Assay | Qiagen | QT00068782 |
| Hs_SNRPE_1_SG QuantiTect Primer Assay | Qiagen | QT00217980 |
| Hs_SNRPF_1_SG QuantiTect Primer Assay | Qiagen | QT01008630 |
| Hs_SNRPG_1_SG QuantiTect Primer Assay | Qiagen | QT00200291 |
| Hs_SEC31A_1_SG QuantiTect Primer Assay | Qiagen | QT00058653 |

#### Primers

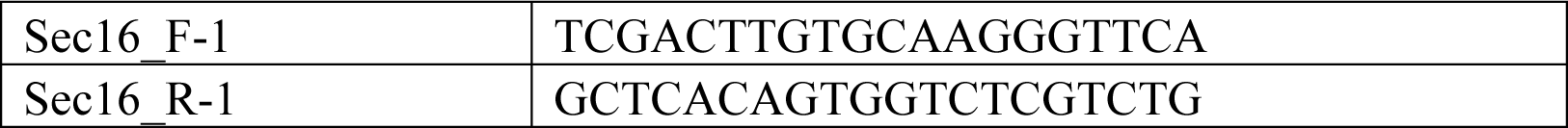
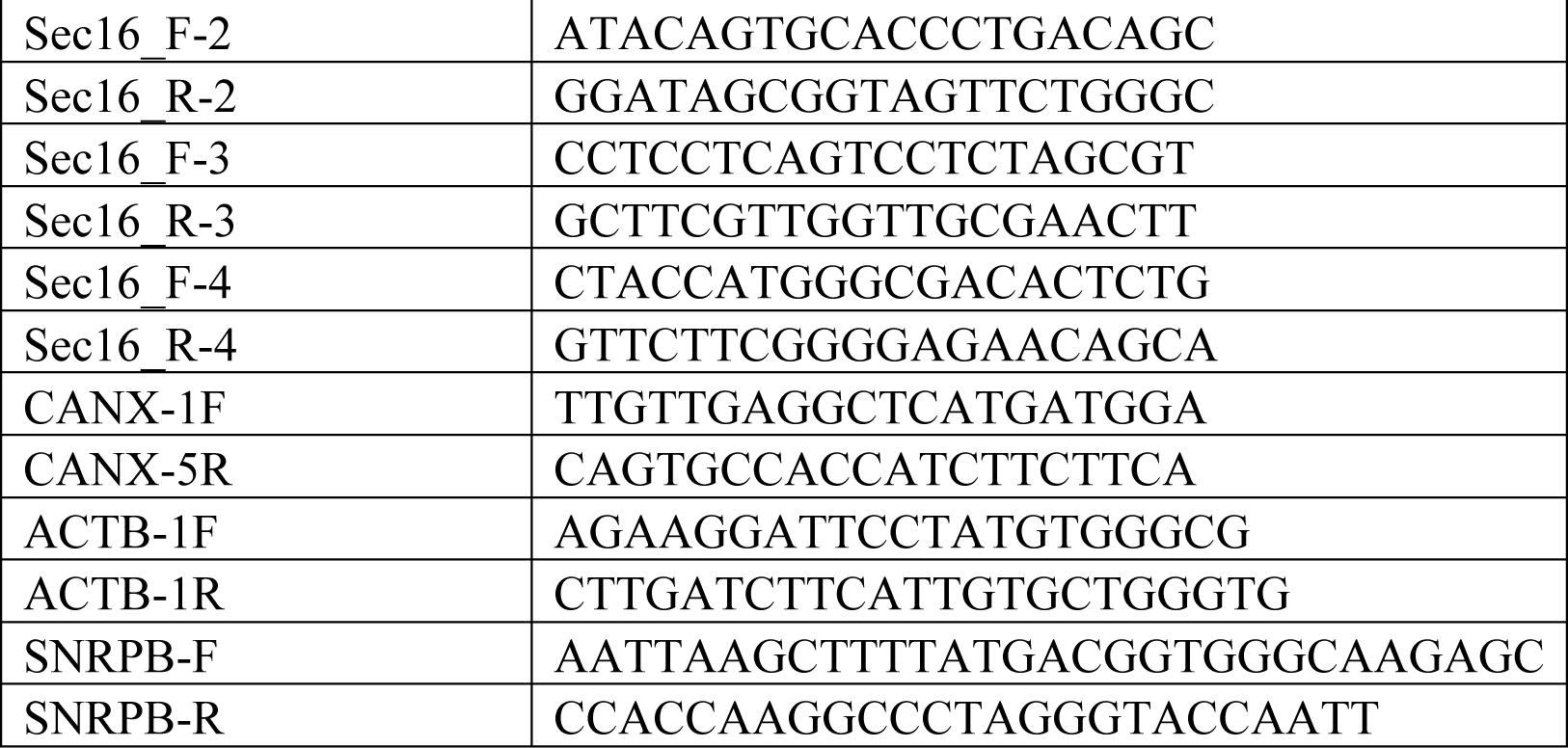

#### siRNAs

|  |
| --- |
| siGenome Smartpool_ATF4 |
| siGenome Smartpool_ATF6 |
| siGenome Smartpool_XBP1 |
| siGenome Smartpool_SNRPB |
| Silencer® Select siRNA_SNRPB2 |
| Silencer® Select siRNA_HS_SNRPB |
| Silencer™ Select Negative Control No. 1 siRNA |

